# A Pangenome-Centralized NLRome Drives Lineage-Specific Diversification and Functional Differentiation in Solanoideae

**DOI:** 10.64898/2026.03.24.712282

**Authors:** Hongliang Zhu, Chao Huo, Lan Wang, Jifen Cao, Zhechao Pan, Zhenchuan Ma, Yan Yuan, Zhijian Zhao

## Abstract

The Solanoideae subfamily comprises numerous economically important crops that deploy a diverse repertoire of nucleotide-binding leucine-rich repeat receptors (NLRs) to defend against pathogens. However, the evolutionary trajectory of these NLR immune receptors remains poorly understood. Here, we dissect NLR family evolution across 23 Solanoideae species by integrating genomic and transcriptomic data. We uncover a distinctive pangenome-centralized architecture wherein a mere 11.56% of soft-core orthogroups contribute 67.81% of the total NLR repertoire, with copy-number variation driven by lineage-specific expansions rather than stochastic distribution or genome size. This finding indicates that Solanoideae plants counteract biotic stress through targeted amplification of specific key families rather than undergoing broad diversification. Phylogenomic analysis resolves NLRs into four subclasses, revealing CC-NLR dominance across all species alongside extreme TIR-NLR degeneration in certain lineages, suggesting profound immune adaptive shifts. Mechanistically, recent NLR expansions are primarily fueled by proximal and tandem duplications, whereas whole-genome duplications contribute in a ploidy- and time-dependent manner, with WGD-derived genes undergoing extensive loss during diploidization. Transcriptional profiling further demonstrates duplication-type-dependent expression patterns, enabling a robust, multilayered defense network upon pathogen infection. Collectively, our study delineates the asymmetric evolutionary trajectory of the Solanoideae NLRome and establishes a framework model that provides new insights into the adaptive evolution of plant immune systems and resistance gene discovery.

## 1. Introduction

In nature, plants are constantly challenged by diverse pathogens, including bacteria, fungi, oomycetes, viruses, and nematodes. Within agricultural systems, these pathogens cause substantial yield losses across major crops. To counter pathogen invasion, plants have evolved a sophisticated innate immune system comprising two primary layers of defense (Jones and Dangl 2006). The first layer, pattern-triggered immunity (PTI), relies on cell surface-localized pattern recognition receptors (PRRs)—primarily receptor-like kinases (RLKs) and receptor-like proteins (RLPs). These PRRs recognize pathogen- or microbe-associated molecular patterns (PAMPs/MAMPs), extracellular effectors, and damage-associated molecular patterns (DAMPs), thereby activating basal defense responses (Zipfel 2014; Ortiz-Morea et al. 2022). The second layer, effector-triggered immunity (ETI), involves intracellular resistance proteins known as nucleotide-binding leucine-rich repeat receptors (NLRs). NLRs directly or indirectly perceive pathogen effectors, triggering a robust immune response often accompanied by localized programmed cell death, termed the hypersensitive response (Cui et al. 2015).

Most plant disease-resistance genes encode NLR receptors, which underpin plant protection against pathogens. Plant genomes typically harbor hundreds of NLR genes, frequently organized in genomic clusters. A canonical NLR protein consists of three domains: a variable N-terminal domain, a central conserved nucleotide-binding (NB-ARC) domain, and a variable C-terminal leucine-rich repeat (LRR) domain. Among these, the NB-ARC domain constitutes the most conserved core structure, while the LRR domain displays high sequence diversity driven by diversifying selection and is primarily responsible for effector recognition (Kourelis and Van Der Hoorn 2018). Based on variations in the N-terminal domain, NLRs can be categorized into four subfamilies: TIR-NLRs (TNLs) containing a Toll/interleukin-1 receptor domain, CC-NLRs (CNLs) with a coiled-coil domain, CCR-NLRs (RNLs) featuring an RPW8-like domain, and the recently identified CCG10-NLRs (Kourelis et al. 2021; Lee et al. 2021; Qureshi et al. 2023). Each NLR subfamily fulfills distinct roles in plant immunity. TNLs and CNLs typically function as pathogen sensors, directly or indirectly recognizing effector proteins. CNLs can assemble into dimeric, tetrameric, pentameric, or hexameric resistosomes that act as cation-selective channels, initiating Ca^2+^ influx and coordinating downstream immune responses. TNLs form resistosomes that function as NAD^+^ hydrolases (NADases), producing secondary signaling messengers such as pRib-AMP/ADP and ADPr-ADP/di-ADPR. In contrast, RNLs lack direct effector recognition capability but instead function as helper NLRs, supporting specific CNLs and TNLs in mediating downstream signal transduction (Wang et al. 2019a, 2019b; Bi et al. 2021; Förderer et al. 2022; Liu et al. 2022; Parker et al. 2022; Liu et al. 2024; Ma et al. 2024).

NLR genes originated from the common ancestor of green plants, with early differentiation into the CNL and TNL subclasses already evident in green algae and bryophytes (Gao et al. 2018; Shao et al. 2019). The RNL group emerged later in gymnosperms, where it functions as a helper NLR alongside TNLs and CNLs (Qureshi et al. 2023; Shao et al. 2016). In angiosperms, NLR genes have undergone two major evolutionary phases: an early period of low copy number lasting until the Cretaceous-Paleogene (K-Pg) event, followed by significant expansion after the K-Pg mass extinction (Li et al. 2010; Shao et al. 2016).

Over long-term evolution, different plant lineages have developed distinct NLR evolutionary patterns under pathogen pressure. For example, Brassicaceae exhibits an expansion-contraction pattern (Zhang et al. 2016), while Poaceae shows a contraction pattern driven by gene loss or frequent deletions (Luo et al. 2012; Shao et al. 2014; Jia et al. 2015), and Cucurbitaceae maintains a relatively low NLR number due to frequent gene loss coupled with limited gene duplication (Wan et al. 2013). Notably, even within the same plant family, species can display markedly different NLR evolutionary patterns, as observed in four orchid species—*Phalaenopsis equestris*, *Dendrobium catenatum*, *Gastrodia elata*, and *Apostasia shenzhenica* (Xue et al. 2020), as well as in Solanoideae species such as tomato, potato, and pepper (Qian et al. 2017).

Solanaceae, an economically and biologically important family of angiosperms, consists of approximately 100 genera and 2700–3000 species widely distributed across temperate and tropical zones. Many of its members hold substantial economic, culinary, medicinal, and ornamental value (Christenhusz and Byng 2016; Raj et al. 2022; Särkinen et al. 2013). Among them, potato (*Solanum tuberosum*), tomato (*Solanum lycopersicum*), eggplant (*Solanum melongena*), pepper (*Capsicum annuum*), and groundcherry (*Physalis* spp.) are important vegetable crops, while medicinal plants such as deadly nightshade (*Atropa belladonna*), henbane (*Hyoscyamus niger*), and jimsonweed (*Datura stramonium*) play important roles in medicine. Furthermore, several Solanoideae species like petunia, tobacco, tomato, and potato are widely used as model organisms in biological research. Solanoideae crops face persistent threats from devastating diseases caused by pathogens such as *Phytophthora infestans*, *P. capsici*, *P. nicotianae*, *Ralstonia solanacearum*, *Alternaria solani*, Tomato spotted wilt virus (TSWV), and Tobacco mosaic virus (TMV), leading to severe agricultural losses. Therefore, an in-depth analysis of NLR genes in Solanaceae is of great significance for understanding of plant immune evolution, mining novel disease-resistance genes, and designing effective resistance-breeding strategies.

Previous studies have shown that NLR genes in Solanaceae species such as tomato, potato, and pepper have undergone significant expansion and contraction across different species, yet a consistent evolutionary trend remains largely absent (Qian et al., 2017). Solanoideae encompasses a variety of economically significant crops, and a cross-species perspective that simultaneously elucidates their NLRome evolution is still lacking. To address this gap, we integrated genomic and transcriptomic data from 23 species representing six genera of Solanoideae to systematically decipher the evolution of the NLRome. Our findings revealed that the Solanoideae NLRome exhibits a “pangenome-centralized” architecture, wherein a few key families are responsible for the majority of NLR copies, accompanied by asymmetric evolution of NLR subclasses (dominance of CC-NLRs and lineage-specific degradation of TIR-NLRs). Furthermore, copy number variations are primarily driven by local duplications such as proximal duplications (PD) and tandem duplications (TD), whereas the contribution of whole-genome duplications (WGDs) is ploidy-dependent and time-dependent. Additionally, NLRs originating from different duplication modes displayed layered expression patterns under pathogen stress, suggesting functional differentiation in immune networks. This study provides critical insights into the evolution and immune response organization of the NLR family in Solanoideae, offering a theoretical basis for resistance gene discovery and disease-resistant breeding.

## 2. Materials and Methods

### 2.1. Mining and Identification of the NLR Gene Family in Solanoideae Clade

Genomic data for 23 species within the Solanoideae subfamily were retrieved from public databases, including the NGDC and the Sol Genomics Network (https://solgenomics.net/) (Supplementary Table S1). Additionally, *Ipomoea batatas*, *Arabidopsis thaliana*, and *Petunia inflata* were selected as outgroup species. All genomes were obtained in annotated format. Candidate NLR genes were identified from the protein-coding reference files of each species using the NLRtracker software (Qureshi et al. 2023). NLRtracker integrates InterProScan analysis and detection of specific core motifs within domains (e.g., P-loop, GLPL) to identify candidate NLR proteins, generating output files containing sequences, annotations, and NB-ARC domain sequences. Each candidate NLR sequence was subsequently validated through clustering and phylogenetic analysis alongside known reference NLRs, allowing manual correction of any misannotations from the automated pipeline.

### 2.2. Phylogenetic Reconstruction of Species and NLR Genes

Orthologous groups were inferred from the protein datasets of all 26 genomes using OrthoFinder (Emms and Kelly 2019). To construct a robust species phylogeny, lineage-specific, low-copy orthologous gene families were refined by selecting families with a copy number <4 and coverage > 90%. The longest copy from each species within these families was selected, and a custom script was applied to remove redundant copies from the orthogroups. Multiple sequence alignment was performed using MUSCLE (Edgar 2004; v5.3), followed by trimming with Gblocksto retain conserved blocks (Castresana 2000). All conserved blocks were concatenated into a supergene alignment, which was used to construct a maximum likelihood species tree with IQ-TREE (Nguyen et al., 2015). For the NLR-specific phylogeny, the conserved NB-ARC domains (>300 bp) extracted by NLRtracker were aligned using MUSCLE. The alignment was refined using MUSCLE, trimmed with trimAl (Capella-Gutiérrez et al. 2009), and a maximum likelihood tree was built with IQ-TREE, employing the best-fit model and 1,000 ultrafast bootstrap replicates for node-support assessment. All phylogenetic trees were visualized and annotated using iTOL (Letunic and Bork 2024).

### 2.3. Gene Loci Visualization and Synteny Analysis

NLR gene loci were mapped and visualized using the output files including GFF3 annotation generated by NLRtracker. BEDtools was used to define gene clusters by intersecting NLR gene coordinates with a defined interval of 30 kb (Quinlan and Hall 2010). The results generated an edited count file, assigning a bin number to each coordinate to create a karyotype file for circos-style visualization using the RCircos package in R (Hao et al., 2020). Synteny analysis was performed based on whole-protein sequence homology. MCScanX, implemented within TBtools (Chen et al. 2023), was used to identify syntenic genomic blocks and homologous gene pairs. The Advanced Circos module in TBtools was subsequently employed to generate intra- and inter-species synteny maps, with NLR genes highlighted and annotated.

### 2.4. Evolutionary Analysis and Gene Duplication Classification

To investigate evolutionary relationships among NLR genes, OrthoVenn2 was used to cluster NLR genes (Xu et al. 2019). Protein sequences of the putative NLR genes from different species were provided with an E-value cutoff of 1e-2 as the default setting to determine common genes across all species. This clustering was complemented by orthogroup inference using OrthoFinder. Subsequently, Gene-family expansion and contraction dynamics were analyzed using CAFE5 (Mendes et al. 2020), with the ultrametric species tree and the OrthoFinder-derived orthogroups as input.

Given the relatively lower genome assembly quality of *Solanum pimpinellifolium*, *Physalis angulata*, *Lycium barbarum*, and *Anisodus luridus*, whole-genome duplication (WGD) analysis was conducted using protein sequences and annotations from the remaining 19 high-quality genomes. For each species, intra-species BLASTP (e-value < 1e-10, top 5 matches) was performed to identify paralogous gene pairs (Camacho et al. 2009). Using *Ipomoea batatas* as the outgroup for all species, the BLAST results and gene position information for each species against *I. batatas* were processed with DupGen_finder to identify potential duplication patterns of NLR genes within the genomes (Qiao et al. 2019).

To assess selective pressures acting on the NLR gene family, the non-synonymous to synonymous substitution rate ratio (Ka/Ks) was calculated to assess whether this gene family primarily underwent purifying selection or diversifying selection during evolution using KaKs_Calculator 3.0 (Zhang 2022). Gene pairs with Ks > 2 were filtered out to avoid the impact of sequence substitution saturation on evolutionary rate estimation.

### 2.5. Expression Profiling of NLR Genes in *Solanum* under Pathogen Challenge

The potato or tomato–*P. infestans* pathosystems represent classic models for studying plant–oomycete interactions. To investigate the role of NLR genes during immune responses, we analyzed their expression patterns in potato and tomato following challenge by *P. infestans*. Publicly available RNA-seq datasets (accessions: PRJNA728373 and PRJNA997093) from infection time-course experiments on potato cultivars Qingshu9 and RH, as well as on tomato (Cai et al. 2025; He et al. 2021) were utilized to elucidate NLR expression patterns and infer their potential roles in disease resistance. The raw sequencing reads were aligned to their respective reference genomes using HISAT2 (Pertea et al. 2016). Transcripts were then assembled and quantified using StringTie (Pertea et al. 2016). Differential expression between infection time points and mock-inoculated controls was processed and evaluated using the Ballgown package (Pertea et al. 2016).

## 3. Results

### 3.1 Subclass -Specificity and asymmetric expansion of the NLRome in Solanoideae

The species phylogeny reconstructed using OrthoFinder was largely consistent with previous studies (Huang et al., 2023). By combining NLRtracker with manual curation, we identified a total of 12,148 NLR genes across 23 Solanoideae species, and characterized the NLR subclass composition and phylogenetic relationships of these species (Fig. 1a). In general, a considerable amount of variation was observed among species in both total NLR counts and subclass proportions, exhibiting clear lineage specificity. NLR copy numbers remained relative stability within Hyoscyameae and Datureae, while greater evident intra-tribal variation was observed in Solaneae, Capsiceae, and Physalideae. It is noteworthy that the cultivated tetraploid potato Qingshu 9 and Cooperation 88, along with *Iochroma cyaneum*, displayed significant NLR expansion. Across all subclasses, CC-NLRs were found to be predominant in every Solanoideae species. TIR-NLRs exhibited pronounced contraction and were even nearly absent in certain lineages such as *Physalis floridana*, displaying a marked trend of degradation. CCG10-NLRs found to be enriched in only a few taxa, while CCR-NLRs exhibited a consistent retention of only 2–5 copies, showing a highly conserved distribution pattern.

**Figure 1.**
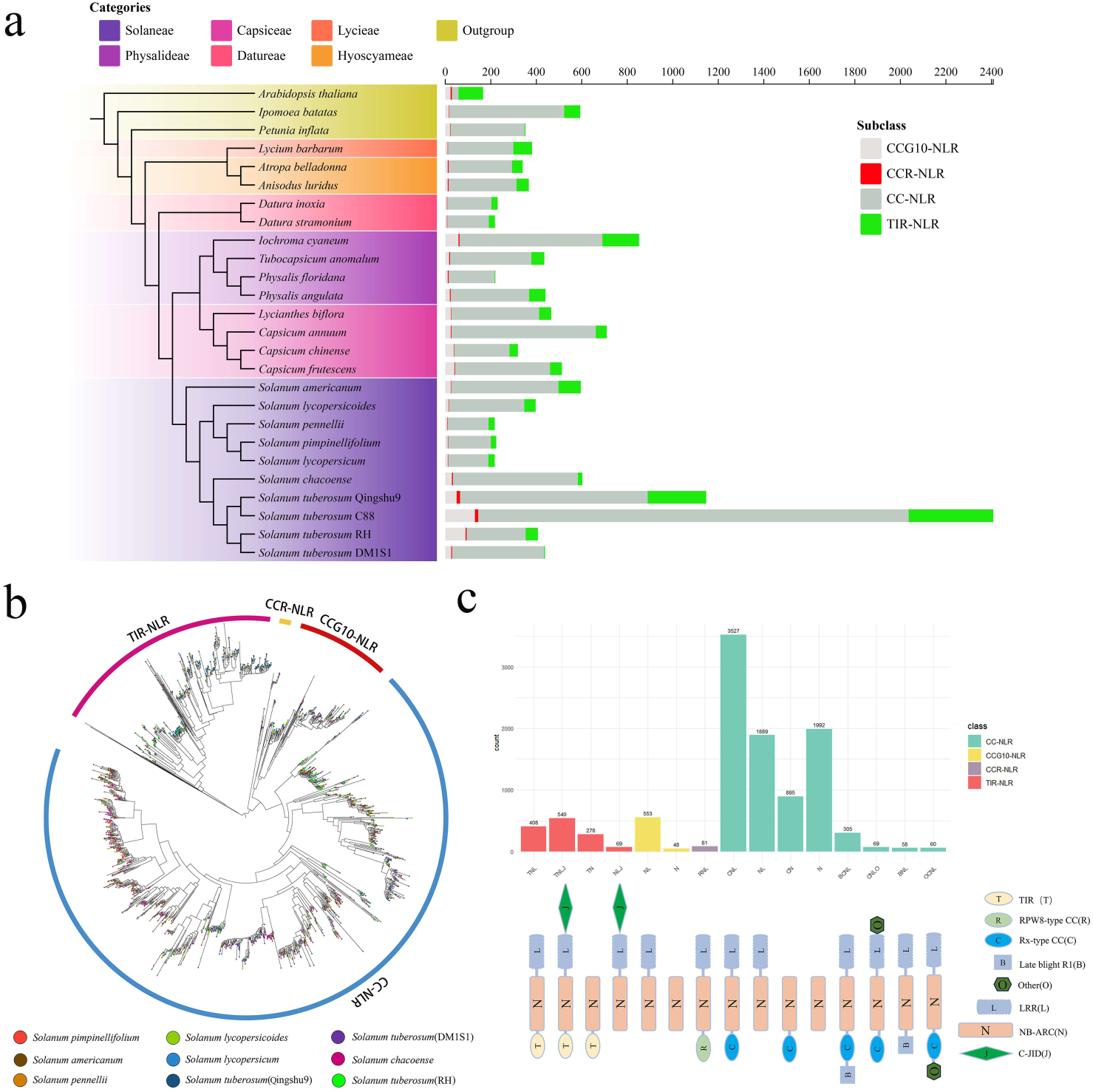
Distribution, phylogeny, and structural diversity of NLR genes in Solanoideae. (a) Bar chart showing the copy numbers of four NLR subclasses (CC-NLR, TIR-NLR, CCG10-NLR, and CCR-NLR) across 23 Solanoideae species, with different colors representing distinct subclasses and clades indicated by colored brackets. (b) Maximu-likelihood phylogenetic tree constructed based on the conserved NB-ARC domains of NLR genes identified in the *Solanum*. (c) Schematic representation of the abundance and domain architectures of major NLR structures within the Solanoideae.

In order to further dissect their evolutionary dynamics, we extracted the conserved NB-ARC domain sequences (>300 bp) from all identified NLRs, yielding 5,954 sequences for phylogenetic reconstruction (Fig. 1b). The resulting phylogeny resolved Solanoideae NLRs into four distinct clades: CC-NLR, TIR-NLR, CCG10-NLR, and CCR-NLR. The CC-NLR clade was found to be overwhelmingly predominant. Within both the CC-NLR and CCG10-NLR clades, we identified lineage-specific expansion subclades primarily composed of members from Capsiceae or Physalideae (Supplementary Fig.1), whereas other clades did not exhibit comparable expansion levels, indicating pronounced asymmetric expansion across different NLR lineages within the Solanoideae NLR family. Integrated protein domain annotation (Fig. 1c) revealed that structural variation among NLRs primarily manifests in the combinatorial patterns of N-terminal recognition domains and C-terminal LRR units. While most genes retained canonical domain combinations, some displayed atypical architectures. In Solanoideae NLRs, CC-NLRs exhibited the highest structural diversity. In the case of TIR-NLRs, beyond their numerical contraction, they also incorporated additional functional domains integration-related variants. Conversely, CCG10-NLRs and CCR-NLRs maintained relatively simple architectures and low copy numbers. Collectively, these findings demonstrate that the Solanoideae NLRome, while preserving core signaling modules, has evolved into a highly plastic immune receptor repertoire through asymmetric expansion, TIR-NLR degradation, and domain recombination.

### 3.2 NLR Abundance is Independent of Genome Size in Solanoideae

NLR gene distribution was visualized by dividing genomes into 30-kb bins and mapping the number of genes present in each bin. The density distribution of NLR genes across the genomes of different species was illustrated in Figure 2. Overall, NLR gene number showed no linear correlation with genome size. For example, *T. anomalum* possesses a large genome (∼5.02 Gb) but contains only 435 NLR genes, presenting a sparse distribution in the density plot (Supplementary Fig. 2). Conversely, the tetraploid potato cultivar Qingshu9 (∼2.67 Gb) and C88 (∼3.05 Gb) harbors 1,147 and 2,408 NLR genes, respectively, exhibiting multiple high-density clusters (Supplementary Fig. 3). Similarly, *I. batatas* (∼837 Mb) contains 594 NLR genes and shows higher NLR density than several larger genomes. Within the Capsicum genus, *C. annuum* and *C. frutescens* have comparable genome sizes but markedly different NLR gene counts, further supporting the decoupling of NLRome size from overall genome size. These findings indicate that NLRome expansion and contraction are primarily driven by evolutionary mechanisms such as gene family-specific amplification, tandem duplication, and gene loss, rather than by genome size alone. Furthermore, the distinct distribution patterns also reflect lineage-specific evolutionary pressures. For instance, the prominent NLR clustering observed in Qingshu9 might be associated with rapid gene family expansion driven by the combined effects of artificial selection and pathogen pressure. Conversely, the sparse NLR distribution in the Datureae clade might result from extensive gene loss and family contraction throughout its evolutionary history.

**Figure 2.**
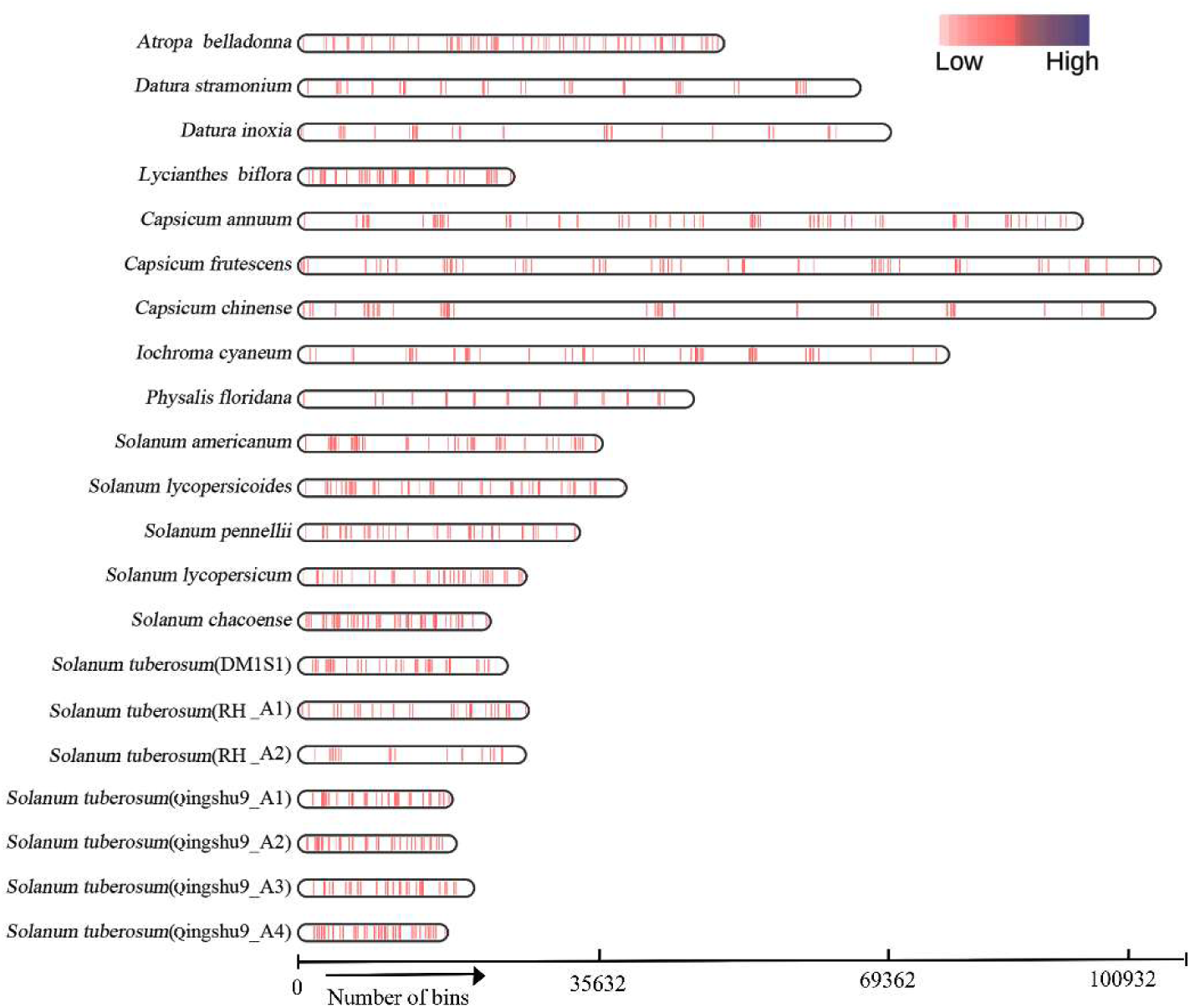
Relationship between NLR gene number and genome size in Solanoideae. Using chromosome-anchored genomes, NLR gene distribution was analyzed in 30-kb sliding windows to assess the relationship between genome size and NLR repertoire organization. For species with haplotype-resolved assemblies, individual haplotypes are distinguished by suffixes (e.g., A1, A2) appended to the species name.

### 3.3 Gene Gain and Loss Dynamics Drive NLR Repertoire Evolution in Solanoideae

The evolutionary dynamics of the NLR gene family were inferred using CAFE5, based on the OrthoFinder-derived species phylogeny. This analysis uncovers distinct branch-specific patterns of gene expansion and contraction among major Solanoideae clades (Figure 3a). The Datureae subfamily underwent an ancestral loss of approximately 17 NLR orthogroups and has continued to exhibit a net contraction trend — a pattern that may reflect specific ecological adaptations. In contrast, the Solaneae subfamily shows high rates of both gene gain and loss, with an overall trend of net loss. This dynamic turnover may reflect efficient purging mechanisms associated with its predominantly selfing reproductive strategy, which could accelerate the removal of nonfunctional or deleterious alleles. The Capsiceae subfamily, however, has undergone sustained and large-scale expansion of gene families throughout its evolutionary history, potentially driven by an elevated molecular evolutionary rate or recurrent tandem duplication events. Within the Physalideae subfamily, *I. cyaneum* has undergone continuous NLR expansion since its ancestral divergence, indicating persistent and strong selective pressures.

**Figure 3.**
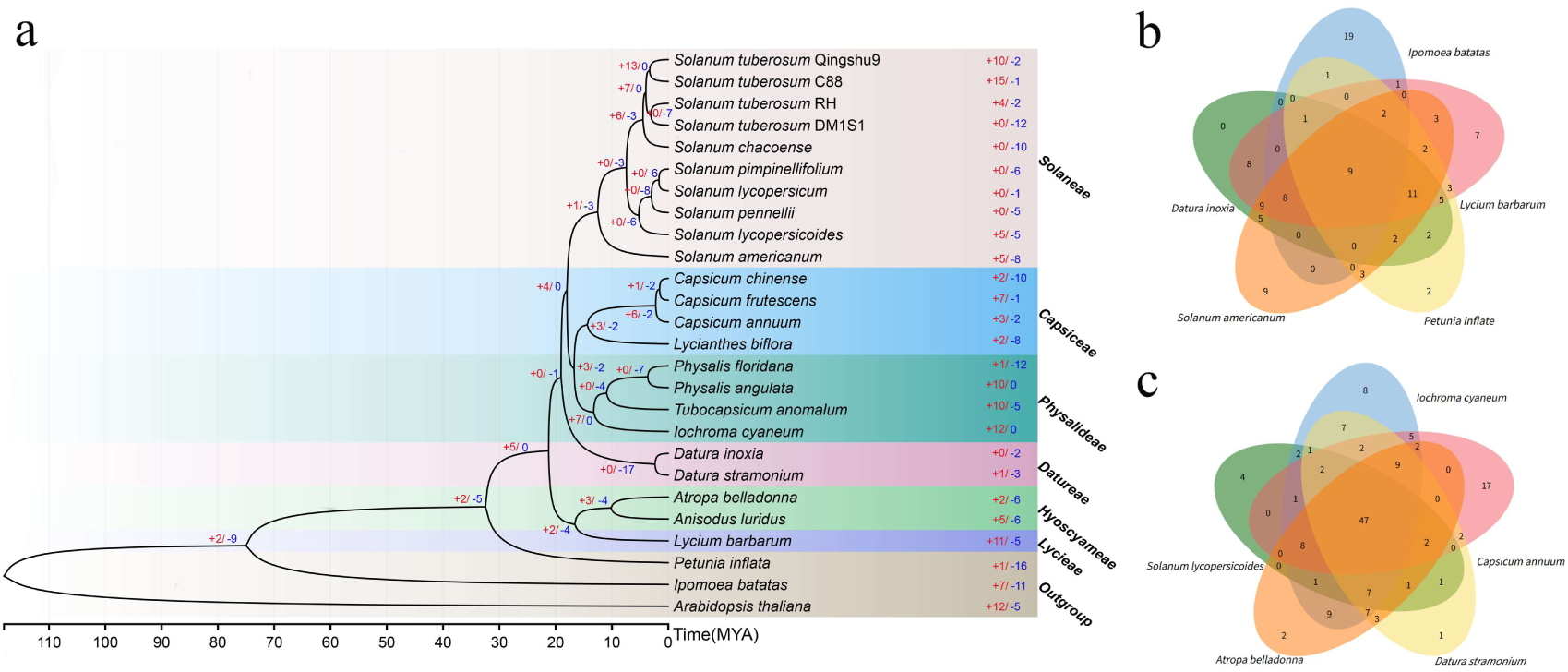
Orthogroup dynamics and gene gain/loss of NLR genes in Solanoideae. **(a)** Phylogenetic tree showing NLR gene count changes across Solanoideae lineages. Node values represent gene family sizes; blue and red numbers indicate gene gains and losses, respectively. **(b, c)** Venn diagrams depicting the distribution of shared and unique NLR orthogroups among representative Solanaceae species.

Venn diagram analyses (Figure 3b, 3c) visually illustrate the distribution of NLR orthogroups across evolutionary depths. These results show that only a limited set of core NLR families is shared across all lineages, while the majority of gene families exhibit pronounced lineage-specific distributions. Together, these markedly divergent patterns of gene gain and loss across Solanoideae reflect the diverse evolutionary strategies these lineages have adopted in response to distinct environmental and pathogenic challenges.

Furthermore, we investigated inter-species chromosomal synteny across Solanoideae (Supplementary Fig. 4). Comparative syntenic analysis revealed that CCR-NLR genes exhibit significantly higher positional conservation across lineages. In contrast, CC-NLR-enriched regions are characterized by extensive rearrangements and cluster remodeling. In *I. cyaneum* which harbors a substantially expanded NLR repertoire, gene expansion appears to be driven by a combination of local amplification and locus translocation mechanisms. This highlights chromosomal-level restructuring as a major driver of lineage-specific NLR diversification.

### 3.4 Independent Evolution and Lineage-Specific Expansion of the NLR Repertoire in Cross-Species Orthogroups

To systematically investigate the evolutionary characteristics of the NLR gene family in Solanoideae, we performed a cross-species orthogroup analysis, clustering all NLR genes into 424 orthogroups (OGs; Supplementary Table S4). This pan-genomic approach provides a more comprehensive representation of genetic variation than a single reference genome.

We first assessed the evolutionary independence of Solanoideae NLRs by comparing them with outgroup species (*Arabidopsis thaliana* from Brassicaceae and *Ipomoea batatas* from Convolvulaceae). Orthogroup clustering revealed fundamental compositional differences between Solanaceae and the outgroups. Core NLR families that underwent massive expansion in Solanaceae (e.g., OG0000000, OG0000001, and OG0000002, each containing ∼600 genes) were rare or entirely absent in the outgroups. Conversely, *A. thaliana* and *I. batatas* also possessed unique expanded families not detected in Solanaceae (e.g., OG0000121 in sweet potato). This strong inter-family exclusivity confirms that the 424 NLR orthogroups in Solanoideae were not retained from an ancient angiosperm ancestor, but rather represent lineage-specific amplification following family-level divergence.

Having established the independent origin of the Solanoideae NLR repertoire, we further dissected the pangenome distribution of these 424 OGs across the 26 studied genomes (Figures 4a, 4b). Statistical analysis classified 9 OGs as ‘Core’ (present in all 26 species), 49 as ‘Soft-core’ (18-25 species), 236 as ‘Dispensable’ (2-17 species), and 130 as ‘Private’ (single species). Although dispensable OGs accounted for 55.7% of all orthogroups, they contributed only 15.9% of total gene copies, suggesting these families are typically small, lineage-restricted, and may represent recent evolutionary innovations or pathogen-specific resistance genes. In contrast, Soft-core OGs, comprising just 11.6% of orthogroups, contributed 67.8% of all NLR genes, indicating that a limited set of conserved families has undergone pronounced copy-number expansion across most species.

**Figure 4.**
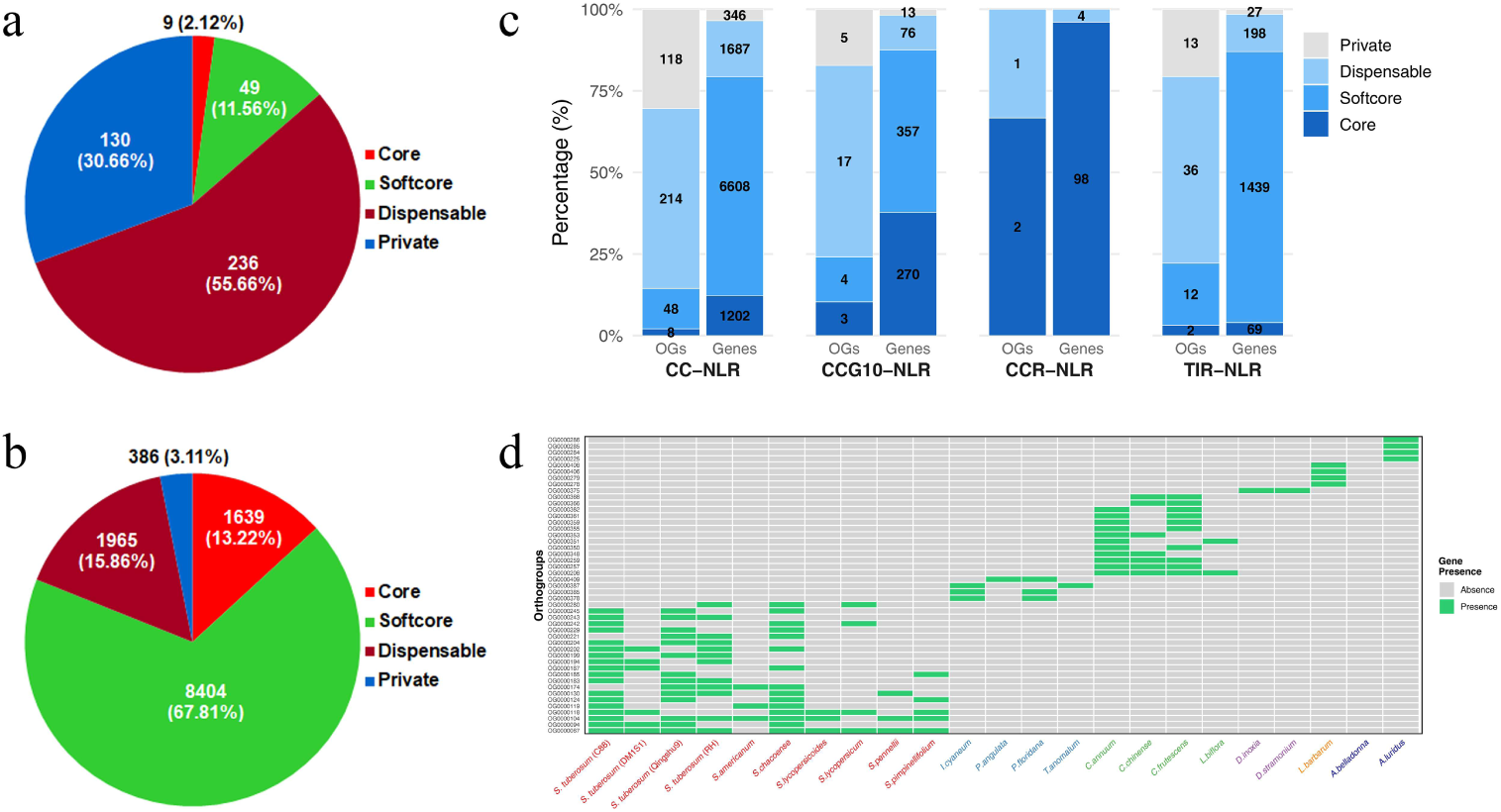
Pangenome analysis of NLR orthogroups across 23 Solanoideae species. **(a)** Proportion of orthogroups (OGs) in each pangenome component (Core, Soft-core, Dispensable, and Private). **(b)** Proportion of total NLR gene counts in each pangenome component. **(c)** Comparison of pangenome components among the four NLR subfamilies (CC-NLR, CCG10-NLR, CCR-NLR, and TIR-NLR). The left bar represents the number of OGs, and the right bar represents the total gene count; numbers on bars indicate raw values. **(d)** Heatmap showing lineage-specific orthogroup distribution (presence criteria: Solaneae ≥ 3 species; Physalideae and Capsiceae ≥ 2 species; Datureae and Hyoscyameae ≥ 1 species; Lycieae ≥ 1 species). The x-axis represents individual species, and the y-axis represents lineage-enriched OGs. Green indicates presence and gray indicates absence.

Classification by NLR subclass revealed distinct evolutionary trajectories (Figure 4c). CCR-NLRs were extremely evolutionary conserved, with 96% of their genes belonging to the core component (represented by only 2 OGs) and showing hardly any lineage-specific expansion. However, CC-NLRs exhibited a notable disparity between family diversity and gene abundance. While 85% of CC-NLR orthogroups (332/388) were dispensable or private, indicating numerous small, lineage-specific families, the Soft-core component, representing only 12.4% of CC-NLR OGs (48/388), contributed 67% of CC-NLR genes (6,608/9,843). This suggests that the CC-NLR subclass primarily relies on large-scale duplication of a few conserved families to maintain a vast foundational gene pool, while continuously diversifying into numerous low-copy, rare families to adapt to changing environments. To identify resistance genes unique to specific plant lineages, we screened for gene families broadly present within a specific lineage but entirely absent in all others. The results show that each major Solanoideae lineage possessed a distinct, clade-specific NLR repertoire (Figure 4d). For instance, both Solaneae (e.g., OG0000067) and Capsiceae (e.g., OG0000208) harbored enriched, lineage-specific gene clusters completely absent in the other lineage, indicating that after divergence, different clades have evolved their own unique combinations of resistance genes to precisely combat pathogen pressures in their respective ecological niches.

In summary, the Solanoideae NLRome exhibits a pronounced centralized architecture (wherein a limited set of soft-core families contribute the majority of NLR copies), alongside the continuous emergence and turnover of lineage-specific gene families. This configuration reflects a layered evolutionary framework comprising a stable core repertoire and a rapidly evolving adaptive repertoire.

### 3.5 Evolutionary Dynamics, Selective Pressures and Duplication Modes in the Solanoideae NLR Family

To investigate the evolutionary dynamics and selective pressures underlying NLR expansion in Solanoideae, we analyzed the Ks distribution, duplication modes: Whole-Genome Duplication (WGD), Tandem Duplication (TD), Proximal Duplication (PD), Transposed Duplication (TRD), and Dispersed Duplication (DSD), and Ka/Ks ratios of paralogous NLR pairs (Fig. 5).

**Figure 5.**
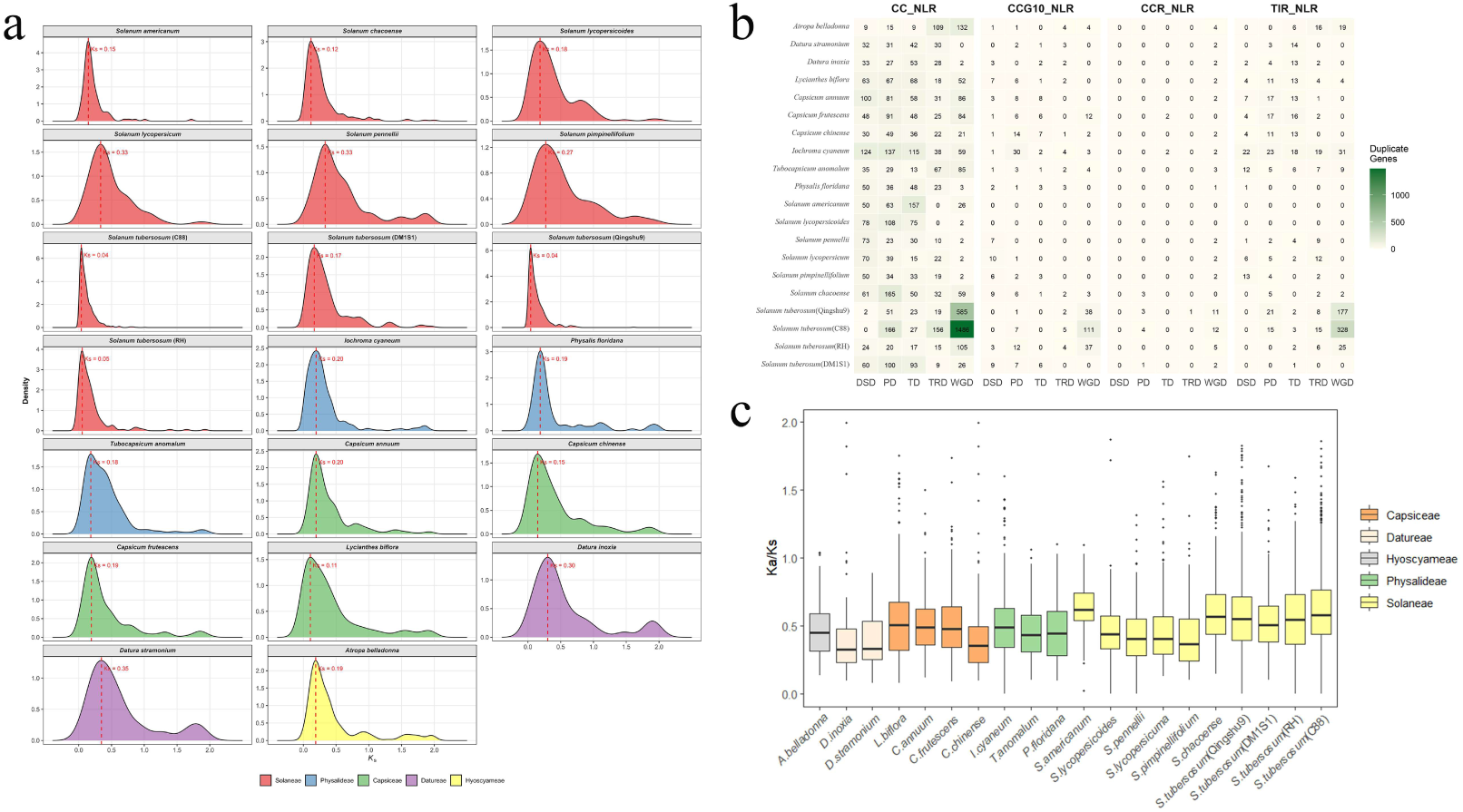
Evolutionary dynamics of NLR genes in Solanoideae. **(a)** Density distribution of Ks values for NLR genes in *Solanum* species. **(b)** Contribution of different duplication mechanisms (WGD, DSD, PD, TRD, and TD) to the expansion of each NLR subclass. Color intensity represents gene count. **(c)** Distribution of Ka/Ks ratios for NLR genes across Solanoideae.

Based on calibrated divergence times (Ks ≈ 0.3), pepper diverged from tomato/potato approximately 19.1 million years ago (Mya) (Choi et al., 2023; Kim et al., 2017, 2014; Tomato Genome Consortium, 2012). Tomato and Etuberosum (potato-like relative) diverged from each other around 14 Mya, followed by a natural hybridization event in the Andean region approximately 8–9 Mya that led to the formation of the tuber-bearing potato lineage (Zhang et al., 2025b). Utilizing this timeline framework, we examined the expansion history of the NLR family. Within the Solaneae clade, distinct Ks peaks were observed among representative genomes. The hybrid tetraploid potato lines Qingshu 9 and Cooperation 88 exhibited very recent peaks (Ks ≈ 0.04), with a high proportion of WGD-derived duplicates, a feature likely attributable to the merger of parental genomes in a hybrid context. In contrast, the doubled haploid potato line DM1S1, having undergone homozygization that eliminated recent heterozygous signals, revealed an earlier expansion peak (Ks = 0.17). The cultivated tomato and its wild relative *S. pennellii* displayed the highest Ks values (0.33, corresponding to ∼21 Mya), suggesting that their major NLR expansion occurred during the tomato–potato divergence. Outside Solaneae, other clades exhibit unique evolutionary patterns with most lineages exhibiting Ks peaks ranging from 0.11 to 0.20, whereas Datureae tended to show more ancient peaks (Ks ≈ 0.30–0.35). Across these diverse species, both TD and PD contributed substantially to NLR retention.

Ka/Ks analysis revealed that the median values across Solanoideae were significantly less than one, thus indicating widespread purifying selection acting on NLR genes. However, potato (Qingshu 9, Cooperation 88) and *S. chacoense* exhibited more dispersed Ka/Ks distributions, characterized by numerous outliers exceeding one. This suggests that subsets of NLRs in these lineages may have experienced episodic positive or diversifying selection, potentially associated with domestication and/or local adaptation. Collectively, these findings demonstrate that NLR expansion in Solanoideae is driven by multiple duplication mechanisms, with recent expansions primarily attributable to local duplications such as TD and PD, whereas the contribution of WGD is both ploidy-dependent and time-dependent.

### 3.6 Duplication-Type-Specific Expression Dynamics Shape a Robust NLR Defense Network in Solanoideae

To systematically characterize the transcriptional regulation of NLR genes, we integrated time-course transcriptomic data from potato and tomato challenged by *Phytophthora infestans* infection. Expression profiles were analyzed across early infection stages (0, 2, 8, 24 h) in diploid potato RH and tomato, and mid-to-late stages (0, 24, 48, 72 h) in tetraploid potato Qingshu9. By overlaying these data with phylogenetic and duplication-type annotations, we constructed a temporal expression atlas of NLR genes across Solanoideae (Figure 6, Supplementary Figure 5, 6).

**Figure 6.**
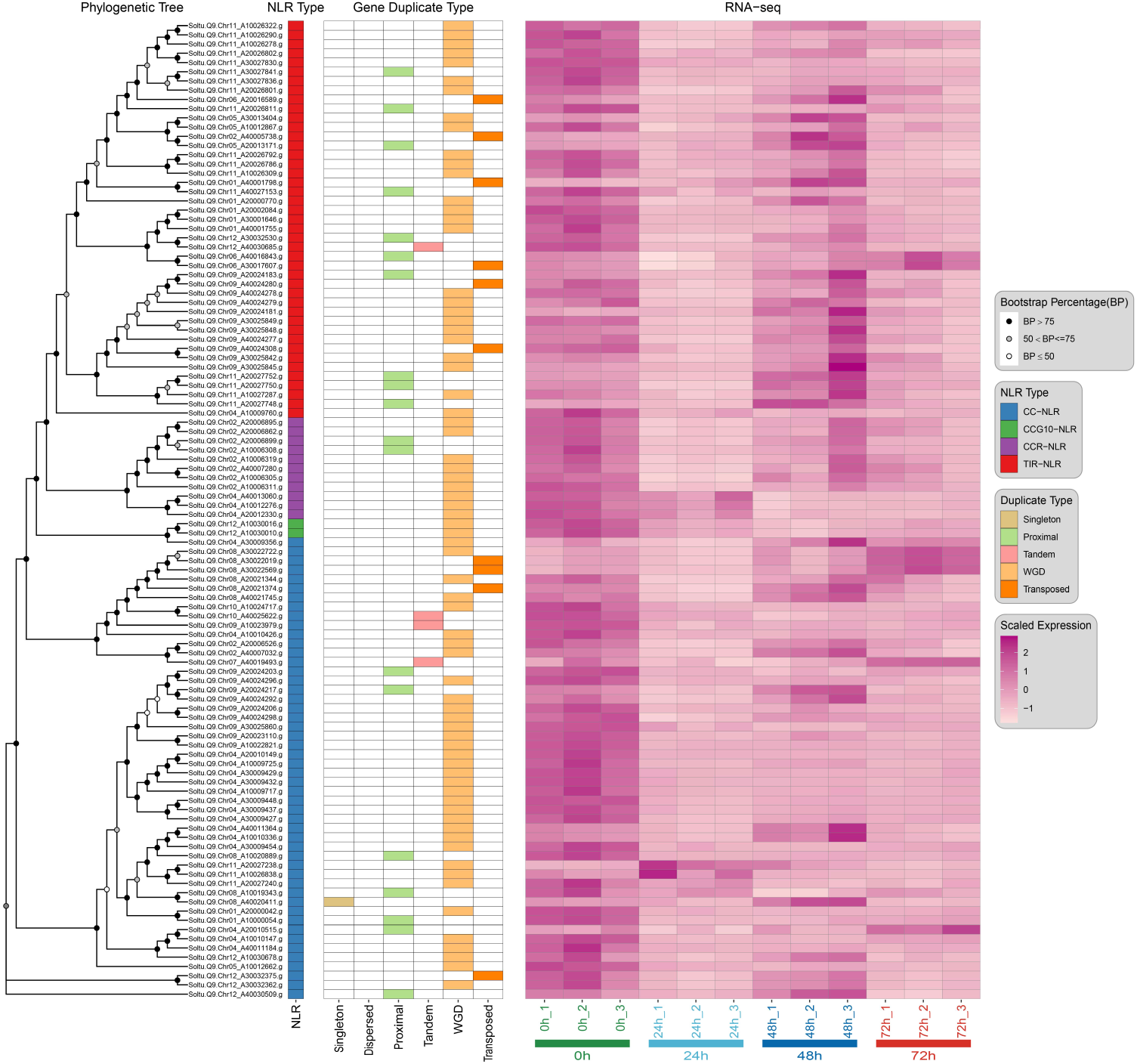
Phylogenetic relationships, duplication types, and temporal expression dynamics of differentially expressed NLR genes. Expression dynamics of NLR genes in *S. tuberosum* (Qingshu9) challenged by *P. infestans* infection. Left: Maximum-likelihood phylogenetic tree of differentially expressed NLRs (node support indicated by bootstrap proportions: BP > 75, 50 < BP ≤ 75, and BP ≤ 50). Middle: Annotations of NLR subclass (CC-NLR, CCR-NLR, TIR-NLR) and duplication mode (Singleton, Dispersed, Proximal, Tandem, WGD, Transposed). Right: RNA-seq expression heatmap at 0, 24, 48, and 72 hours post-inoculation (hpi); three biological replicates are shown per time point.

First, we analyzed the expression trends of different NLR subclasses. The CC-NLRs, which contain multiple key NRC members, emerged as the most active subclass and served as the primary contributors to transcriptional upregulation in response to pathogen infection. Collectively, the CC-NLRs exhibited significant upregulation induction upon biotic stress and played the central role in immune activation. In contrast, although phylogenetically contracted in Solanaceae, the TIR-NLR subclass maintains a relatively high basal expression level in the absence of pathogen stress. However, it rapidly and dramatically downregulated during early infection (0-2h post-inoculation). This suggests that these ancient TIR-NLRs might contribute to maintaining basal immune homeostasis but are transcriptionally vulnerable to suppression by virulent pathogens.

CCR-NLR genes demonstrated remarkable evolutionary and expression conservation across species, exhibiting extremely high expression in the uninfected state but rapidly declining after infection. For example, RHC02H1G2728.2 in potato RH and Soltu.Q9.Chr02_A20006899.g in Qingshu9 are orthologous genes, as are RHC02H2G2563.2 with Soltu.Q9.Chr02_A10006308.g, and RHC02H2G2560.2 with Soltu.Q9.Chr02_A10006305.g. These three gene pairs show remarkable consistency across different genetic backgrounds: under healthy, unchallenged conditions (0h), their expression is very high; however, after *P. infestans* inoculation, their expression dropped rapidly, remaining low in Qingshu9 until 24h post-inoculation. This phenomenon strongly confirms that, under normal conditions, these core CCR-NLRs are crucial and are likely prime targets for pathogen attack. These genes are readily perturbed in early infection, leading to temporary suppression of their expression. Fewer differentially expressed genes were identified in the CCG10-NLR subclass, which was predominantly present in potato RH, showing an overall trend of upregulation.

Integrating expression data with duplication modes revealed a strong association between a gene’s duplication history and its transcriptional regulation pattern. Genes originating from ancient Whole-Genome Duplication (WGD) events primarily exhibit constitutive expression and are prone to suppression. Under normal growth conditions, these genes maintain very high expression abundance across all tissues. However, in the earlier infection stage (0-2 hours), their transcription did not increase further but instead remained low, as confirmed by Qingshu9 data, persisting until 24 hours post-inoculation. This suggests that WGD-derived NLRs function primarily in maintaining basal transcriptional levels but are transcriptionally fragile during early pathogen attack. Conversely, genes originating from non-WGD mechanisms (e.g., tandem duplication) generally exhibit inducible expression profiles. These genes showed minimal expression in uninfected plants but were rapidly and strongly induced within 2-8 h post-infection(hpi), with some peaking during mid-to-late infection stages (48-72 h). This pattern indicates that recently expanded, non-WGD NLR clusters have evolved pathogen-responsive regulatory circuits that enable rapid transcriptional activation, suggesting that new gene clusters generated by non-WGD mechanisms tend to evolve transcriptional response mechanisms highly sensitive to pathogen stimuli.

Among the differentially expressed NLRs, we identified sequences with high homology to multiple classic Solanaceae R genes, including *RPP13*-like, *RGA*, *R1A*/*R1B*, *RPP2A*, and NRC-related members. The majority of upregulated genes originated from non-WGD duplication events and predominantly belonged to soft-core orthogroups (Supplementary Table 7). Collectively, despite some heterogeneity among individual candidates, a convergent trend was observed wherein non-WGD expansion, infection-induced expression, and homology to classical resistance determinants co-occurred in most genes, suggesting that they collectively represent a prioritized set of pathogen-responsive resistance candidates.

Integration of duplication origin with transcriptional response revealed a systematic link between NLR replication history and expression behavior: anciently retained copies tended toward basal expression, whereas recently expanded non-WGD members predominantly exhibited pathogen-induced expression. This points to a layered functional division of NLRs at the transcriptional level according to their duplication origin.

## 4. Discussion

Solanoideae crops, including tomato, potato, pepper, eggplant, and tobacco, represent a vital component of global agriculture but are susceptible to numerous devastating pathogens, resulting in substantial yield losses (Foster and Hausbeck 2010; Fry 2008). Elucidating the molecular basis of disease resistance and breeding resistant varieties in Solanoideae species is therefore crucial for their agricultural sustainability. Through systematic identification and analysis of NLR genes across 23 representative Solanoideae species, this study reveals the complex and dynamic evolutionary landscape of the immune receptor repertoire in this subfamily.

We observed significant interspecific variation in NLR copy numbers (ranging from 217 to 2,408) that was independent of genome size, consistent with findings in other plant families such as Brassicaceae, Fabaceae and Arecaceae (Zhang et al. 2016; Li et al. 2021; Qureshi et al. 2023). A recent study in potato also reported NLR numbers ranging from 348 to 1105 with no correlation to haploid genome size (Wang et al. 2025). Together, these results indicate that NLRome size is shaped primarily by lineage-specific expansion and contraction events rather than by passive scaling with genome size. In Solanoideae, NLR exhibit an overall contraction pattern in Datureae and Hyoscyameae, possibly reflecting diploidization after ancient polyploidization. Conversely, Solaneae, Capsiceae, and Physalideae lineages show a general trend of continuous expansion, possibly reflecting the more complex pathogen pressures faced by these groups. Pangenome analysis further highlighted a centralized architecture: although soft-core NLR families represent only 11.56% of orthogroups, they contribute 67.81% of all NLR genes, indicating that immune adaptation in Solanoideae relies heavily on selective amplification of a limited set of key NLR families.

Whole-genome duplication (WGD) generates a large number of gene copies, a subset of which are retained and can facilitate genetic innovation through subsequent functional or expression differentiation (Arrigo and Barker 2012; Maere and Van de Peer 2010; Mayrose et al. 2011). Genetic and functional changes in retained gene duplicates may enable organisms to exploit new ecological opportunities or cope with new environmental challenges (Fawcett et al. 2012; Schranz et al. 2012). Solanaceae has undergone three recent WGD events, including a whole-genome triplication (WGT) prior to the divergence of the family (∼81 million years ago, Mya) and two subsequent WGDs approximately 37 and 20 Mya (Huang et al. 2023). Despite these ancient events, our analysis indicates that extant NLR genes are predominantly driven by dispersed duplication (DSD) and tandem/proximal duplication (TD/PD). For instance, Datureae retains ancient tandem duplication clusters, while the tomato lineage primarily relies on DSD.

Notably, the substantial number of WGD-type genes (Ks=0.04) observed in hybrid tetraploid potatoes (Qingshu9 and Cooperation 88) does not represent ancient WGD remnants but rather reflects genome complexity arising from recent hybridization and artificial breeding processes, involving subgenome fusion and allelic differentiation. This discrepancy reveals an important evolutionary principle that NLR genes derived from ancient WGDs likely underwent significant loss during diploidization (Zhang et al. 2025a). Consequently, the main body of the modern NLR repertoire consists of genes generated later by other duplication mechanisms like DSD and TD, which exhibit higher long-term retention rates. Furthermore, synteny analysis revealed a lack of NLR homology between Solanoideae and outgroups, further confirming that the NLR repertoire of Solanaceae was not inherited from the common angiosperm ancestor but was remodeled after family-level divergence through frequent interchromosomal rearrangements and in-situ amplification. Synteny analysis further supports the lineage-specific origin of Solanoideae NLRs.

Different NLR subtypes exhibit markedly distinct evolutionary trajectories in Solanoideae. This differentiation pattern reflects deeper-level restructuring of the plant immune system in both architecture and function. CC-NLRs constitute the predominant proportion in all Solanoideae species. Phylogenetic analysis reveals they have formed multiple lineage-specific expanded clades, particularly within Capsiceae and Physalideae. This may be related to specific pathogen pressures faced by these lineages (e.g., *Phytophthora capsici* in pepper, certain leaf spot diseases in *Physalis*). In stark contrast to the widespread expansion of CC-NLRs, TIR-NLRs show a continuous degradation trend within Solanoideae, being nearly completely lost in species like *P. floridana* and *S. tuberosum* (DM1S1). The remarkable contraction of TIR-NLRs in certain Solanoideae lineages may reflect shifts in immune architecture across different phylogenetic groups. On the one hand, numerous CC-NLR-type resistance genes have been cloned and widely applied in Solanaceae crops, conferring resistance against diverse pathogens, such as viruses, oomycetes, and nematodes etc. Representative examples include Sw-5b against TSWV in tomato (Tong et al., 2023), Mi-1.2 against root-knot nematodes in tomato (Adil et al., 2025), and *Rpi-blb1*, *Rpi-vint1.1*, *Rpi-amr1*, *Rpi-amr3*, *Rpi-amr4*, *Rpi-amr16*, *Rpi-amr17*, *Rpi-brk1*, and *Rpi-cph1* against *P*. *infestans* in potato (Lin et al., 2023; Wang et al., 2025). These cases further reinforce the notion that the CNL-mediated ETI represents a central pillar of the immune system within the Solanaceae. On the other hand, TNL-mediated immunity typically depends on co-evolved signaling modules such as EDS1–SAG101–NRG1. If core components of this pathway are missing or subject to regulatory constraints in certain lineages, the TNL system may become particularly prone to degradation (Liu et al., 2021). Furthermore, the remaining TIR-NLRs may have acquired functional diversification through structural domain integration (e.g., the C-terminal jelly-roll/Ig-like domain, C-JID), thereby being retained in specific lineages (Ma et al., 2025). CCR-NLRs, functioning as helper NLRs, are maintained at a low and conserved copy number (2-5) across all Solanoideae species. CCG10-NLRs are a recently defined subtype, but their copy number and distribution vary significantly among different lineages, being relatively abundant in Capsiceae but scarce in Datureae and Hyoscyameae. This distribution pattern likely reflects differential evolution of various Solanaceae lineages under selective pressures from pathogens.

Expression profiling further revealed the profound influence of gene evolutionary history on NLR transcriptional regulation patterns. Our study found that NLR genes exhibit two distinct patterns in response to pathogen infection. The majority of ancient CCR-NLRs and WGD-derived genes maintain relatively high expression levels under normal growth conditions but show a rapid decrease in transcription during early infection. This phenomenon suggests that while these core genes are responsible for maintaining basal immune surveillance, their transcriptional stability is vulnerable to disturbance and suppression during the early stages of pathogen invasion. In contrast, NLR genes recently amplified via non-WGD mechanisms such as tandem duplication exhibit marked inducible characteristics: they have extremely low expression under normal conditions but are rapidly and substantially upregulated upon infection. This indicates that such newly generated genes, produced through recent duplication, are primarily employed to launch specific defense responses upon infection. This complementary pattern—where ancient genes are prone to suppression while newly evolved genes are rapidly induced—reflects an adaptive strategy formed during the long-term evolution of the plant immune system. It involves a division of labor among genes of different origins to construct a more robust defense network. Our results demonstrate that NLR expression during infection is structured by duplication history: CC-NLRs drive induced immune responses, while ancient WGD-derived NLRs are susceptible to early transcriptional suppression. This duplication-type-specific expression partitioning enables Solanoideae plants to maintain both stable baseline immunity and dynamically inducible defense, forming a layered and robust NLR-mediated immune network.

We propose an integrated model for the evolution and functional differentiation of the NLR family in Solanoideae (Fig. 7). This model reveals a distinctive “pangenome-centralized” architecture of the Solanoideae NLRome, characterized by the decoupling of NLR expansion from genome size and its non-random, pronounced concentration within a limited set of core families. Within this architecture, a small number of soft-core orthogroups harbor the majority of NLR copies. Different NLR subclasses display markedly asymmetric expansion patterns: CC-NLRs predominate and have undergone pronounced lineage-specific amplification; TIR-NLRs show an overall trend of contraction; and CCR-NLRs are highly conserved and primarily associated with WGD-derived duplicates, reflecting significant asymmetric differentiation. Furthermore, local duplication modes such as TD and PD constitute the primary drivers of family expansion, whereas WGD events are largely linked to the long-term retention of a subset of conserved members. Consistent with this observation, NLRs originating from different duplication modes exhibit divergent expression strategies: non-WGD expanded members tend to be inducible upon infection and may constitute a rapid response repertoire; conversely, WGD-associated/conserved NLRs generally maintain higher basal expression but are often suppressed during early infection stages, potentially contributing to basal surveillance and immune homeostasis. It should be noted, however, that this model is primarily inferred from pangenomic and transcriptional analyses, and the underlying molecular mechanisms await further functional validation.

**Figure 7.**
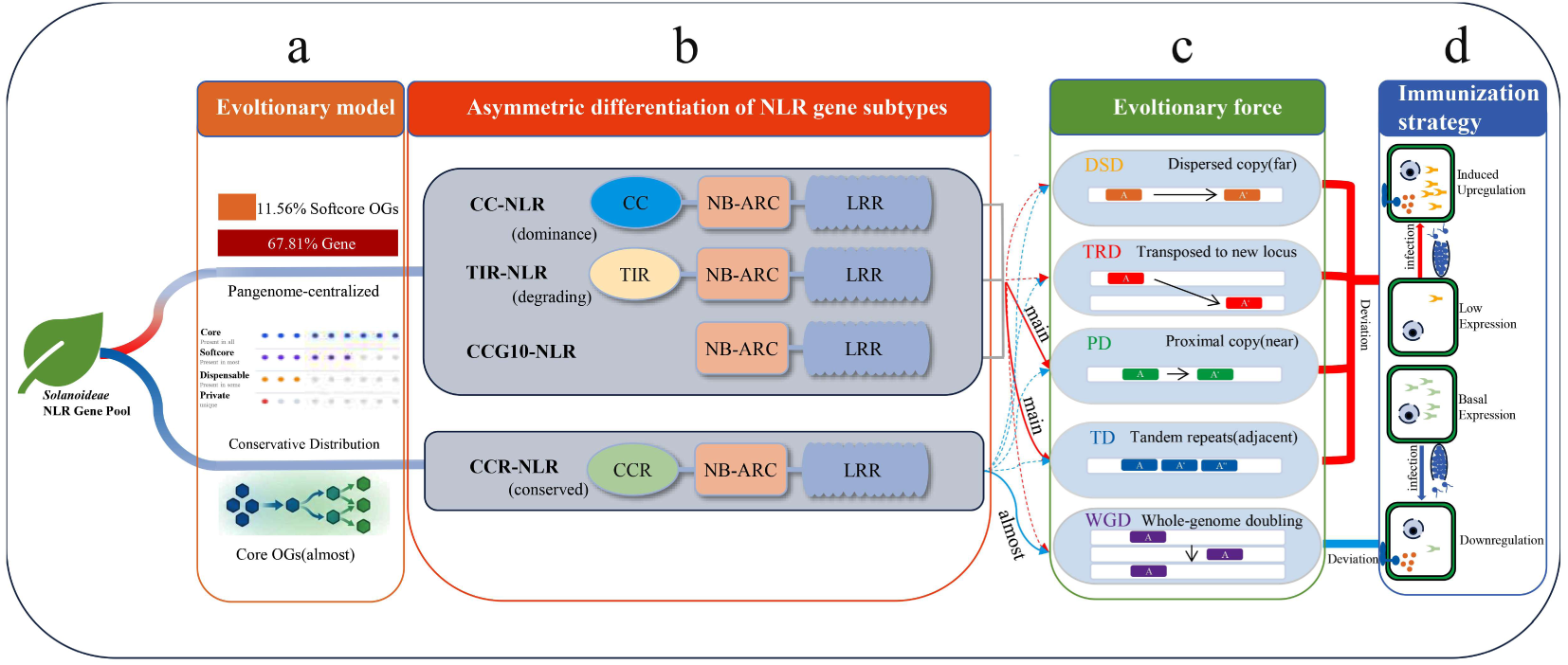
The framework model of dynamic evolutionary trajectory of the Solanoideae NLRome. **(a)** Schematic depicting the pangenome-centralized architecture, wherein a limited number of soft-core orthogroups harbor the majority of NLR copies across species. **(b)** Asymmetric evolution of NLR subclasses, showing the predominance of CC-NLRs, the degenerative trend of TIR-NLRs, and the conserved nature of CCR-NLRs. **(c)** Differential contributions of duplication mechanisms to NLR expansion, with TD and PD serving as primary drivers of recent amplification, whereas WGD events predominantly contribute to the retention of a small set of conserved CCR-NLRs. **(d)** Summary of immune strategies associated with distinct duplication origins: non-WGD expanded NLRs tend to be pathogen-inducible, whereas WGD-derived NLRs generally maintain higher basal expression but are often suppressed during early infection.

In summary, this study not only systematically delineates the dynamic evolutionary trajectory of the NLR family in Solanoideae. Furthermore, it highlights the central role of lineage-specific expansion and duplication-type-dependent expression division in shaping plant immunity. These findings provide new insights into the adaptive evolution of plant immune systems and an important theoretical foundation for future disease-resistance breeding in Solanoideae crops, particularly for harnessing specific NLR resources from wild relatives.

## Acknowledgements

This work was supported by grants from the National Natural Science Foundation of China (nos. 32460050 and 32360226), the Yunnan Seed Laboratory Project (202205AR070001-9) and Joint Special Project of Agricultural Basic Research of Yunnan (202401BD070001-094).

## Author contributions

H.Z., Y.Y., and Z.Z. designed the study. H.Z., C.H., L.W., and Z.Z. performed the study. H.Z., C.H., L.W., J.C., Z.P., Z.M., Y.Y., and Z.Z. analyzed data; H.Z. and Z.Z. wrote the manuscript. All authors read and approved the manuscript.

---

**Figure S1.**
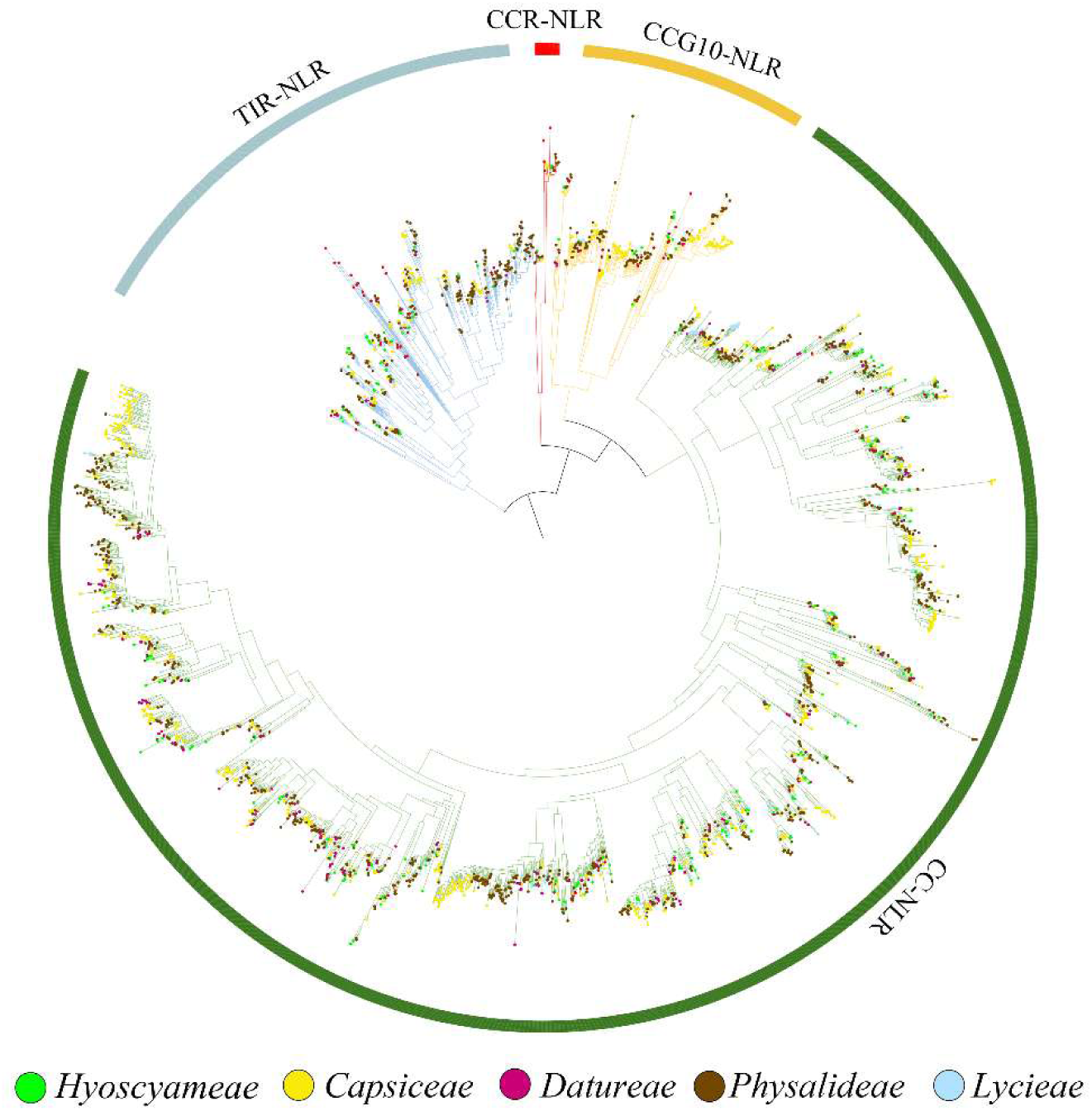
Phylogenetic tree of NLR genes from other Solanoideae species.

**Figure S2.**
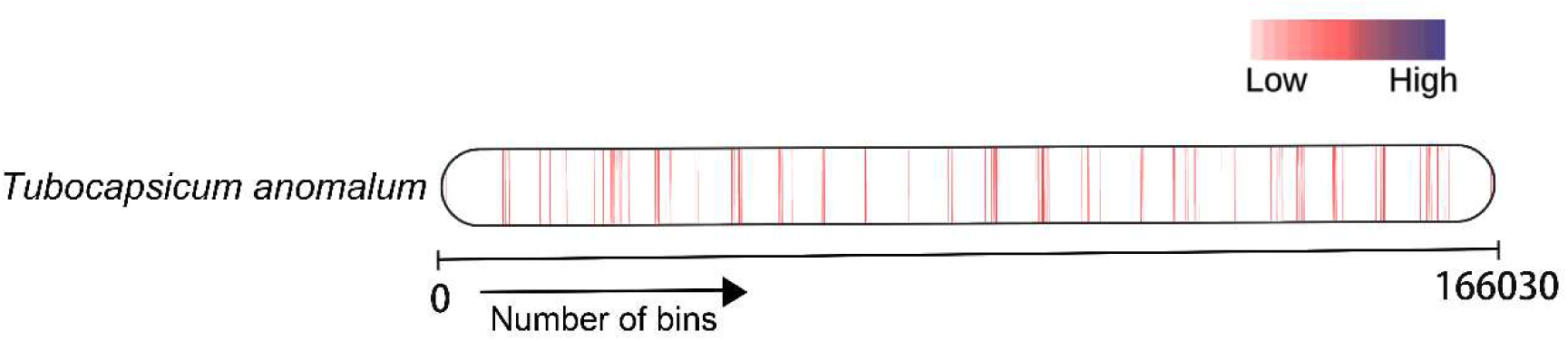
NLR gene density map of *Tubocapsicum anomalum*.

**Figure S3.**
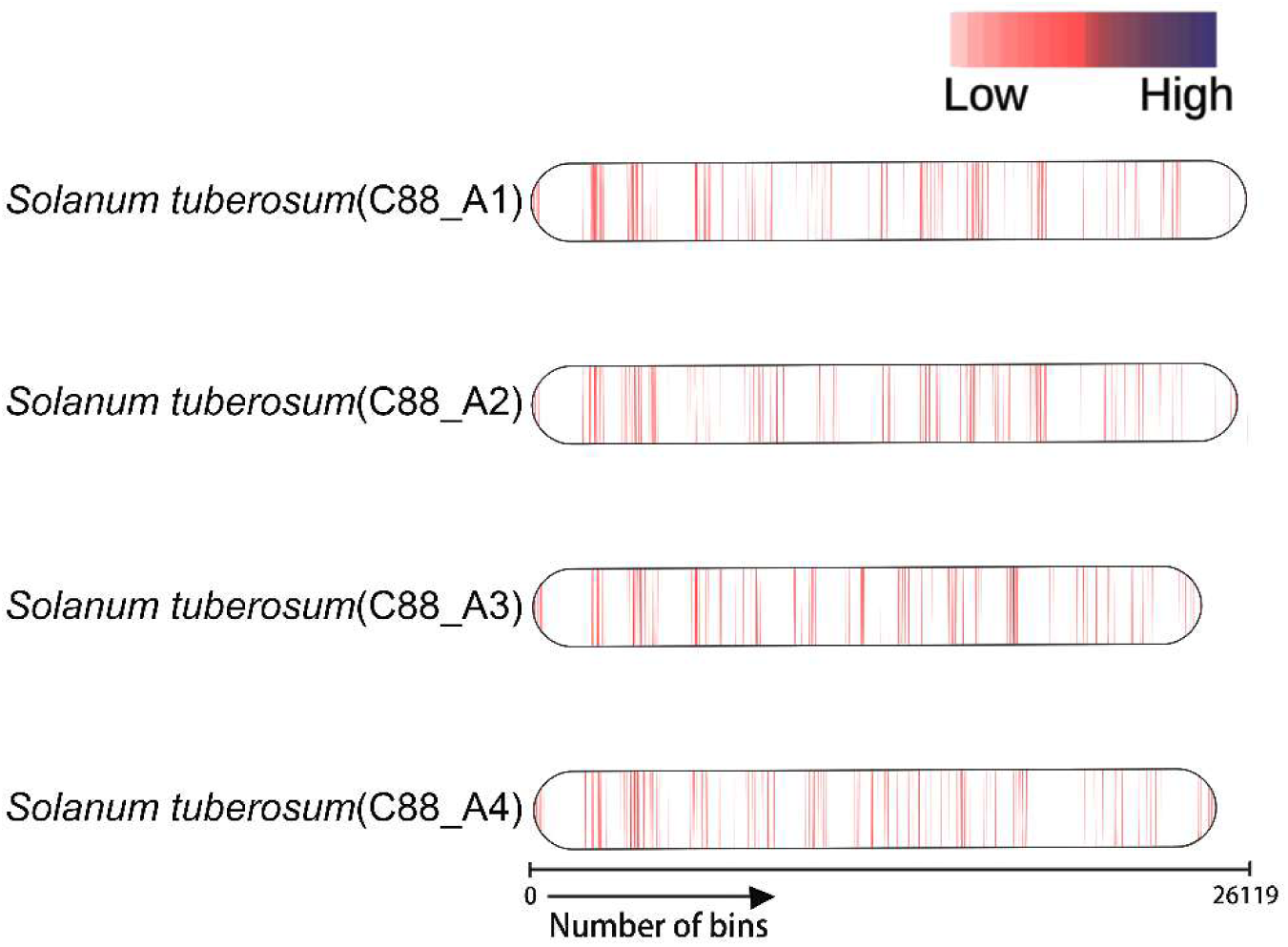
NLR gene density map of *Solanum tuberosum* (Cooperation 88).

**Figure S4.**
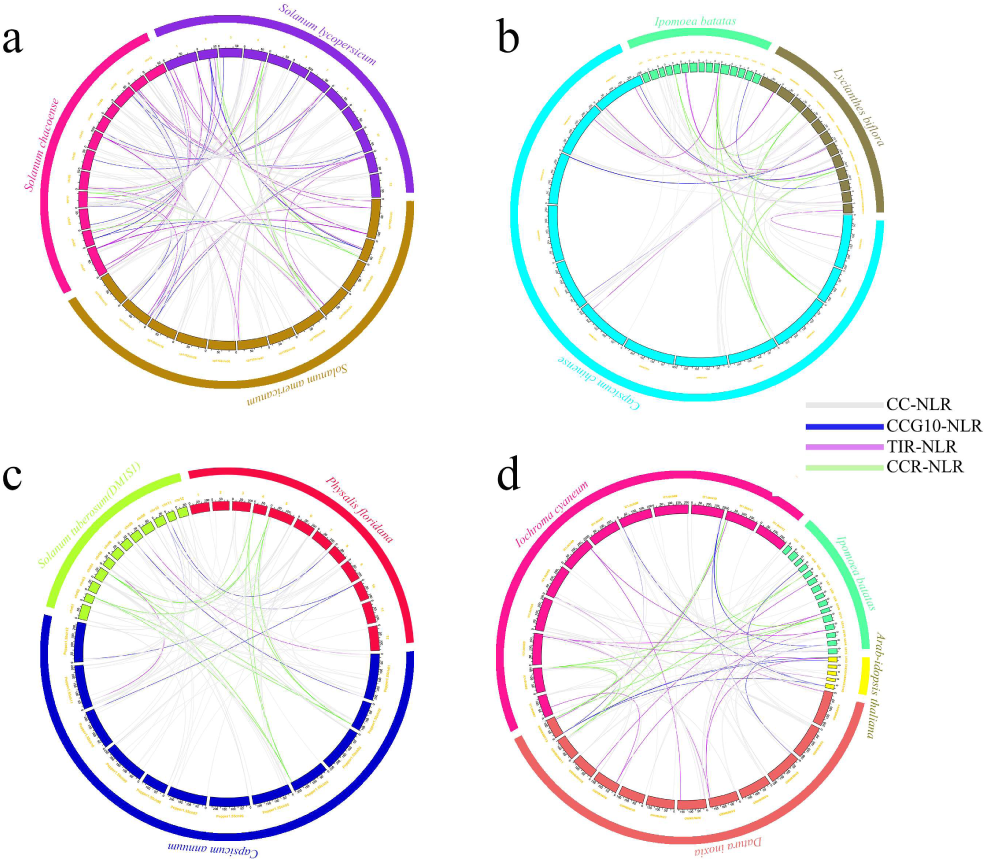
Inter-species synteny analysis across Solanoideae. (a) Genome-wide syntenic relationships within the Solaneae lineage: comparative analysis of *S. americanum*, *S. chacoense*, and *S. lycopersicum*. (b) Syntenic analysis of Capsiceae species with divergent NLR copy numbers and genome characteristics: *C. chinense*, *L. biflora*, and *I. batatas*. (c) Syntenic relationships among *P. floridana*, *C. annuum*, and *S. tuberosum* (DM1S1). (d) Syntenic relationships between Solanoideae and outgroup species: *I. cyaneum*, *D. inoxia*, *A. thaliana*, and *I. batatas*.

**Figure S5.**
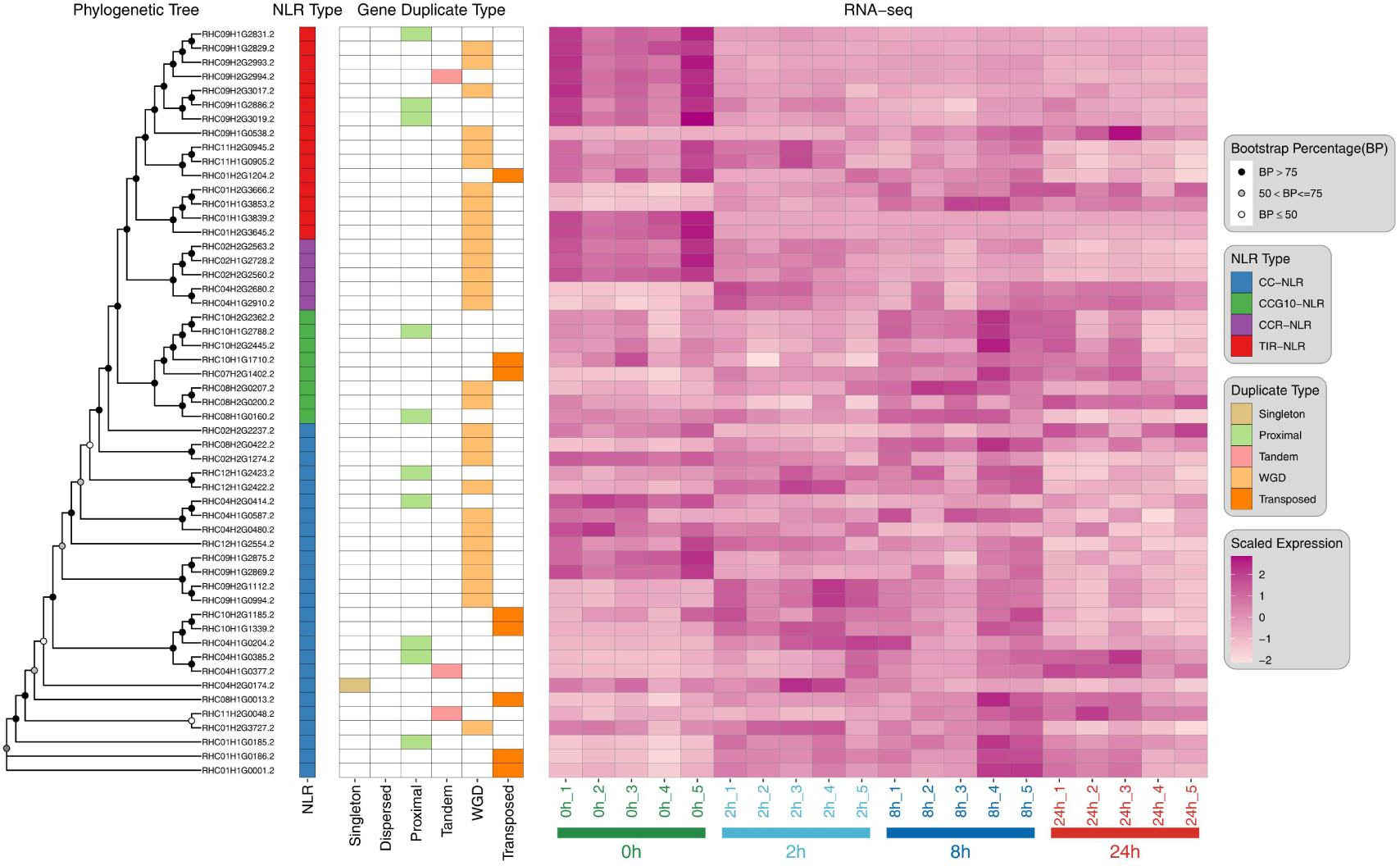
Phylogenetic relationships, duplication types, and temporal expression dynamics of differentially expressed NLR genes in diploid potato (RH).

**Figure S6.**
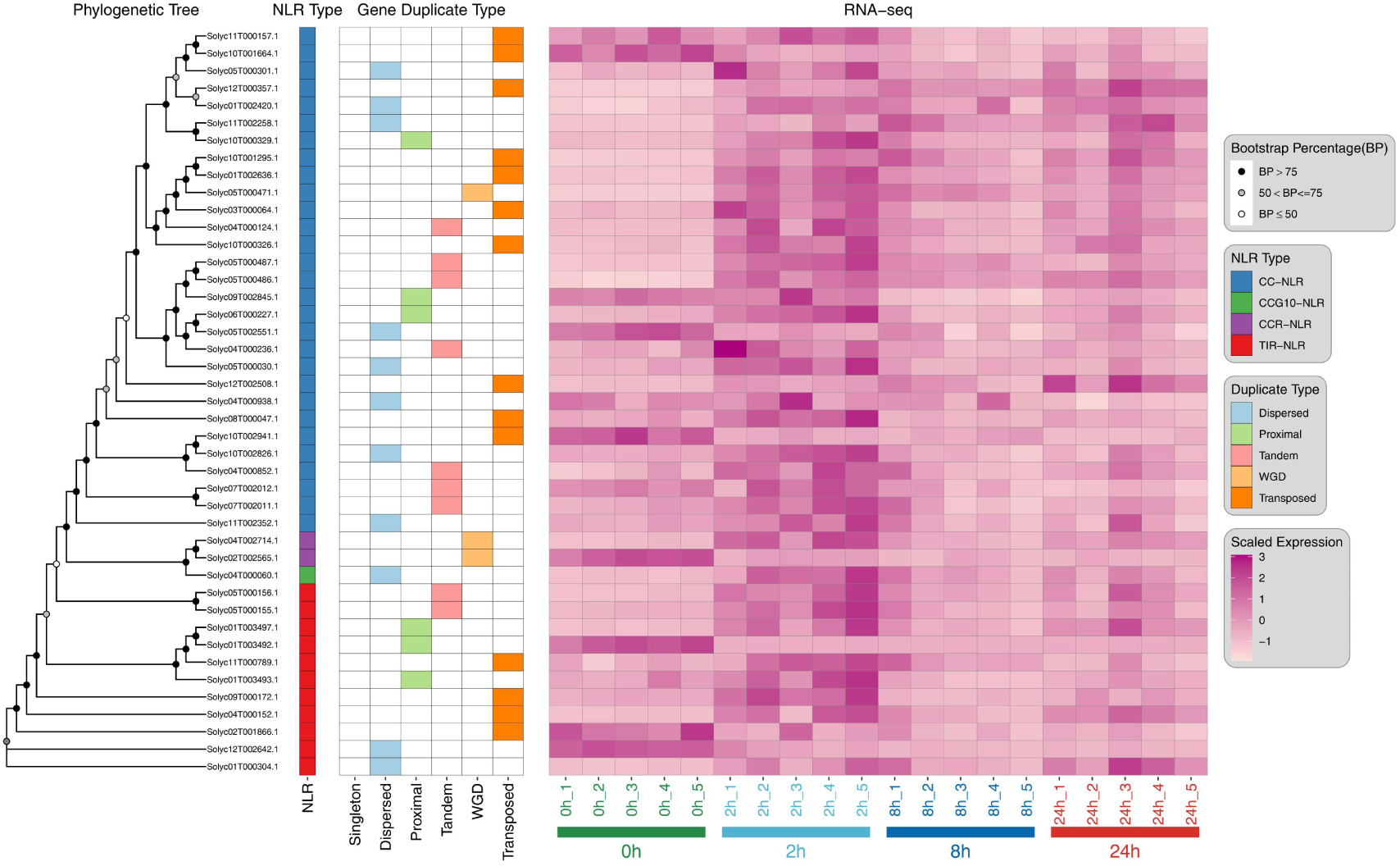
Phylogenetic relationships, duplication types, and temporal expression dynamics of differentially expressed NLR genes in tomato.

